# Subjective time is predicted by local and early visual processing

**DOI:** 10.1101/2020.10.30.362038

**Authors:** Yelena Tonoyan, Michele Fornaciai, Brent Parsons, Domenica Bueti

**Author notes:** correspondence to Yelena Tonoyan and Michele Fornaciai.

## Abstract

Time is as pervasive as it is elusive to study, and how the brain keeps track of millisecond time is still unclear. Here we studied the mechanisms underlying duration perception by looking for a neural signature of subjective time distortion induced by motion adaptation. We recorded electroencephalographic signals in human participants while they were asked to discriminate the duration of visual stimuli after translational motion adaptation. Our results show that distortions of subjective time can be predicted by the amplitude of the N200 event-related potential and by the activity in the Beta band frequency spectrum. Both effects were observed from occipital electrodes contralateral to the adapted stimulus. Finally, a multivariate decoding analysis highlights the impact of motion adaptation throughout the visual stream. Overall, our findings show the crucial involvement of local and low-level perceptual processes in generating a subjective sense of time.

## INTRODUCTION

Time is pervasive in all human activities, and our sense of time is foundational to our very existence. For instance, knowing the precise time at which to reach out and grasp a prey or to cross a busy street require an accurate estimate of time’s passage. Despite such foundational importance, and the growing amount of studies in recent years on this topic, how the brain processes and represents temporal information is still unclear. Several accounts of time perception have been proposed in the past decades, involving different brain mechanisms like a centralized pacemaker-accumulator (Treisman et al., 1990), detectors of oscillatory activity in the striatum (Meck & Benson, 2002), or the intrinsic dynamic of sensory neurons (Buonomano & Maass, 2009). Although the pacemaker-accumulator remains the most popular model – as it can account for many properties of time perception – none of these theoretical frameworks has yet reached general consensus. Conversely, evidence showing different properties of time perception at different time scales suggests that multiple overlapping time-keeping mechanisms may exist in the brain (Fornaciai, Markouli, & Di Luca, 2018; Karmarkar & Buonomano, 2007; Spencer, Karmarkar, & Ivry, 2009; Wiener, Matell, & Coslett, 2011).

Crucial to reaching a deeper understanding of the mechanisms of time perception is to assess how physical temporal information is translated into a *subjective* sense of time. Indeed, perceived time often deviates from physical time, showing that temporal processing is very malleable and prone to distortions. For instance, the perceived duration of a stimulus is strongly modulated by its physical properties, like motion (Brown, 1995; Kanai et al., 2006), size (Xuan et al., 2007), numerosity (Javadi & Aichelburg, 2012; Togoli et al., 2020), and the context in which the stimulus is presented (Fornaciai et al., 2018; Karmarkar & Buonomano, 2007; Kristjánsson, Vuilleumier, Schwartz, Macaluso, & Driver, 2007; Spencer et al., 2009). Understanding the neural pathway leading to a subjective sense of time and the mechanisms responsible for temporal distortions is thus essential to reach a comprehensive and mechanistic understanding of time perception.

In the present study, we leverage on the temporal distortions induced by motion adaptation to assess the brain processes linked to perceived time. Indeed, it has been shown that adaptation to fast motion strongly reduces the apparent duration of a subsequent stimulus (Ayhan, Bruno, Nishida, & Johnston, 2009; Bruno, Ayhan, & Johnston, 2010; Burr, Tozzi, & Morrone, 2007; Fornaciai, Arrighi, & Burr, 2016; Johnston, Arnold, & Nishida, 2006; Latimer, Curran, & Benton, 2014) – an effect that has been named “duration compression.” While the properties of this adaptation effect are debated – especially concerning whether it occurs in a retinotopic (Bruno et al., 2010; Latimer et al., 2014) or spatiotopic (Burr, Cicchini, Arrighi, & Morrone, 2011; Burr, Tozzi, & Morrone, 2007) reference frame – it nevertheless offers a powerful tool to understand the brain mechanisms underlying our subjective sense of time.

To assess the neural signature of subjective time, we asked participants to perform a duration discrimination task (i.e., comparing the duration of a constant reference stimulus to that of a variable comparison duration) during electroencephalographic (EEG) recording. Crucially, in different conditions, participants were adapted to a fast (20 deg/s) or a slow (5 deg/s) moving stimulus, delivered to the same retinotopic or spatiotopic (i.e., screen coordinates) coordinates of the duration stimulus (i.e., the reference stimulus), or in a neutral location.

The idea was to look at how these different adaptation conditions affected the duration perception of a physically identical stimulus (i.e., the reference) and to identify a neural signature of these differences in perception. To preview, event-related potentials (ERPs) results at the level of the N200 component – a component linked to motion processing (Hoffmann et al., 2001) – time locked to the onset of reference stimulus and recorded from electrodes contralateral to the reference presentation, could reliably predict the distortion of perceived time induced by adaptation. Moreover, changes in Beta band frequency observed most prominently in occipital electrodes during adaptation offset and in the first half of the reference stimulus presentation, successfully predict perceptual distortions. Finally, a multivariate decoding analysis shows that motion adaptation impacts the visual pathway at different latencies from reference onset. These results show that subjective time is linked to processes happening locally and relatively early in the visual processing stream.

## RESULTS

The effect of motion adaptation on perceived time was tested with a behavioural paradigm in which we asked healthy human participants (N = 32) to discriminate the duration of pairs of visual stimuli after undergoing motion adaptation. The stimuli used in the duration discrimination task were arrays of moving dots, moving coherently from left to right at a speed of 10 deg/sec (for more details see *Materials and Methods*). The reference stimulus of fixed duration (500 ms) was presented in the upper visual quadrant, the variable comparison duration (ranging from 300 to 700 ms) in the lower quadrant, both stimuli were centred on the vertical midline with a vertical (centre-to-centre) eccentricity of 5 degrees of visual angle from the middle of the screen. In the adaptation phase, prior to duration discrimination, participants were presented with a similar visual stimulus, from now on called the “adaptor,” whose spatial position, speed, and motion direction relative to the reference stimulus was manipulated. Based on previous studies (Bruno et al., 2013a; Burr et al., 2007; Fornaciai et al., 2016; Johnston et al., 2006b) showing changes in perceived duration induced by motion adaptation depending on speed, motion direction and spatial position of the adaptor stimulus with the respect to the reference, we decided to test the effect of motion adaptation in five different conditions. These conditions were: (1) 20 deg/s adaptation in which the adaptor and reference were in the same *retinotopic* coordinates (i.e., corresponding to the projected retinal position of the adaptor after an eye movement of 10 deg away from the first fixation point; 20R); (2) 20 deg/s adaptation in *spatiotopic* coordinates, in which adaptor and reference were in the same spatiotopic position (i.e., same position in screen coordinates; 20S); (3) 5 deg/s spatiotopic adaptation (05S); (4) 20 deg/s spatiotopic adaptation, but with adaptor and reference moving in opposite directions (i.e., *inverse* motion condition; 20I); (5) 20 deg/s adaptation in a neutral location, that was neither the retinotopic nor the spatiotopic position of the reference stimulus (i.e., *no-topic*; 20N). Besides the adaptation condition, duration discrimination performance was also measured in a baseline condition without adaptation (*unadapted* condition). A depiction of the experimental procedure is shown in Fig. 1A.

**FIGURE 1.**
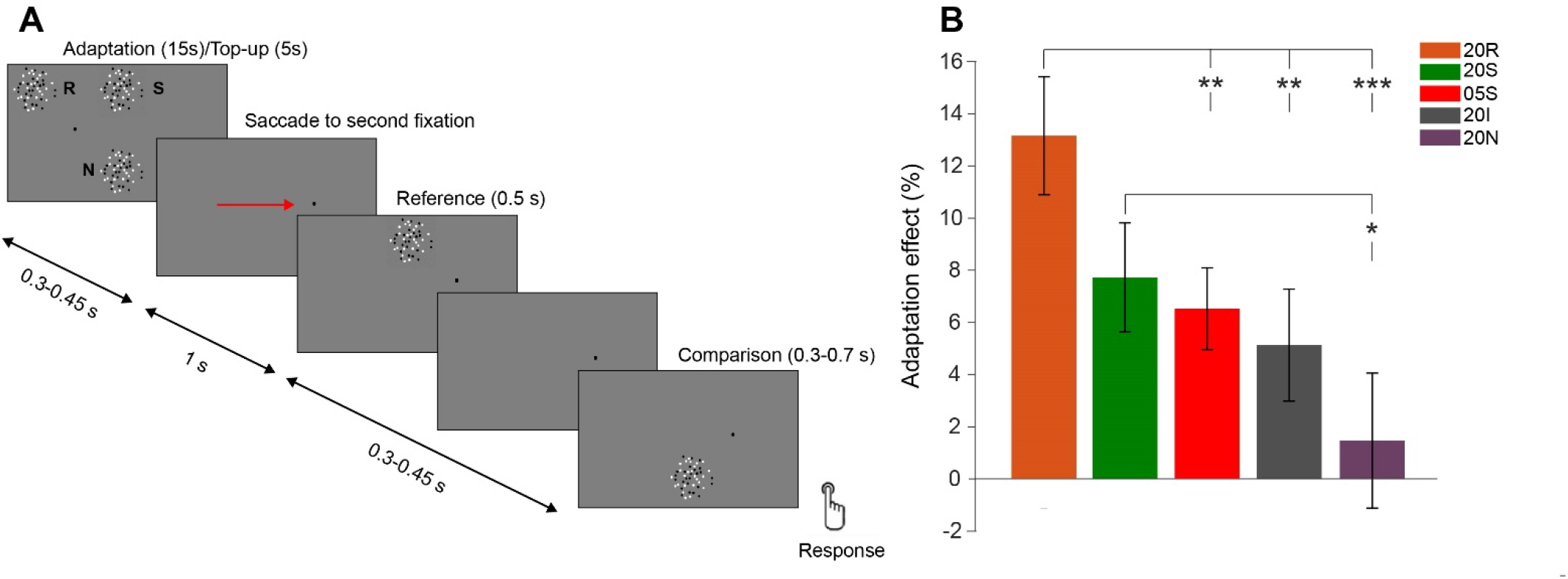
Experimental procedure and behavioural results. (A) Schematic representation of the experimental paradigm. The experimental procedure involved first an adaptation phase (15 s in the first trial, 5 s top-up in the following trials) in which a motion adaptor stimulus was presented in one of three possible positions, corresponding to the spatiotopic (S) or retinotopic (R) coordinates of the reference after an eye movement, or neither of these two reference frames (i.e., no-topic; N). The adaptor could also move at different speeds (i.e., 5 deg/s or 20 deg/s) and in the same or opposite direction compared to the reference stimulus. After the adaptation or top-up phase, the first fixation point (on the left part of the screen) was replaced by a second fixation point on the opposite side (right), 10 degrees away from the first one, cueing the participant to make a saccade on the new fixation. The reference stimulus was presented after the second fixation onset, and was followed by the presentation of the comparison. After the comparison, the fixation point turned red, and participants were instructed to indicate whether the reference or the comparison stimulus lasted longer. After providing a response, the next trial started automatically after 1.3-1.45 s. (B) Behavioural results. Adaptation effects across the different adaptation conditions. Effects are reported in terms of normalized difference of each condition against the unadapted condition, and represent the difference in the reference perceived duration caused by the different types of motion adaptation. Error bars are SEM. Stars refer to the significance level of paired sample t-tests between conditions, performed considering a corrected α value of 0.0125. * p < 0.0125, ** p < 0.01, *** p < 0.001. The legend codes correspond to different adaptation conditions, i.e., 05S = 5 deg/s spatiotopic adaptation, 20I = 20 deg/s inverse motion spatiotopic adaptation, 20S = 20 deg/s spatiotopic adaptation and 20R = 20 deg/s retinotopic adaptation.

As a behavioural measure of the effect of motion adaptation on perceived duration, we focused on changes in the point of subjective equality (PSE; i.e., the duration of the comparison stimulus that is judged 50% of the times longer than the reference) across the different adaptation conditions. To achieve a measure of the net effect of adaptation on duration estimates, we calculated an adaptation effect index. This was the normalized difference in PSE between each adaptation condition and the unadapted baseline condition (see *Materials and Methods* for more details). Fig. 1B shows the results of the behavioural analysis. To assess the effect of adaptation on the reference perception we ran a one-way repeated measures ANOVA on the adaptation index. The analysis showed a significant modulation of adaptation conditions on participants perception (F(4,31) = 6.21, p < 0.001, η^2^ = 0.443). To explore further these modulations we performed a series of one-sample t-tests against the null hypothesis of zero adaptation effect (Bonferroni-corrected α value equal to 0.01). The results showed a significant effect in the 20S (t(31) = -3.64, p < 0.001, Cohen’s *d* = 0.91, average adaptation effect = 7.56%), 20R (t(31) = - 5.7, p < 0.001, *d* =1.43, average effect = 12,86%), and 05S (t(31) = 4.07, p < 0.001, *d* = 1.02, average effect = 6.37%) adaptation conditions. No significant effect was observed instead in the 20I (t(31) = -2.35, p = 0.02, *d* = 0.58, average effect = 5.02%), and 20N (20N; t(31) = -0.5, p = 0.58, *d* = 0.14, average effect = 1.42%) conditions. We further performed a series of paired t-tests comparing all the individual conditions against each other, for this series of comparisons, we considered a Bonferroni-corrected α = 0.0125 (i.e., considering the 4 tests needed to compare a given condition to all the others). The results of these tests across the different adaptation conditions (Fig. 1B) suggest that the most robust duration compression effects were observed in the 20R and 20S adaptation conditions. Namely, the effect in the 20R condition resulted to be significantly different from the 05S, 20I, and 20N condition (p = 0.007, 0.0012, and < 0.001, respectively), while the effect in the 20S condition was significantly different compared to the 20N (p = 0.01) condition. On the other hand, no other condition yielded effects significantly different from the control 20N condition (all p > 0.0125). No significant difference was observed between the 20S and 20R conditions (p = 0.02). The magnitude of adaptation effects at the individual level is shown in Fig. S1 (see *Supplementary Online Materials*).

At the neural level our main goal was to explore the neural responses during the presentation of the reference stimulus – a physically identical stimulus that according to the preceding adaptation condition was perceived differently.

To assess the effect of adaptation at the neural level we looked at event-related potentials (ERP), changes in power across difference frequency bands, and we performed a multivariate decoding analysis. All these analyses focused on the reference stimulus presentation. The only exception was the analysis in the frequency domain, where we deemed important to check for possible entrainment effects during adaptation and in the transition from adaptation to reference stimulus presentation.

We started analysing the event-related potentials (ERP) time-locked to the onset of the adapted reference stimulus, in a set of posterior occipito-parietal channels of interest (O1, Oz, O2, PO3, PO2). The average ERPs across the set of channels of interest, corresponding to the different adaptation conditions, are shown in Fig. 2A. Although the difference is relatively small, the brainwaves plotted in Fig. 2A appear to be sorted according to the strength of the observed behavioural effects, especially at around 200 ms after stimulus onset, corresponding to the N200 component. Namely, the larger the duration underestimation (i.e., shift in average PSE), the smaller the negative amplitude of the N200. A one-way repeated measure ANOVA indeed showed a main effect of adaptation condition on the N200 amplitude (averaged across a time window spanning 140-240 ms), suggesting that brain activity at the level of this component is modulated by the different adaptation conditions (F (4,31) = 4.14, p = 0.001, η^2^ = 0.73). The scalp topography of N200 displayed in Fig. 2B shows that after all the adaptation conditions the response evoked by the reference tends to peak at electrodes contralateral to its position on the screen. The only exception to this pattern is the no-topic and the unadapted conditions, showing much less marked lateralization compared to the other conditions. This can be explained by the lack of duration compression in these adaptation conditions, suggesting again a relationship between the behavioural effect of adaptation and the activity in the N200 latency window. To better assess the relation between the N200 component and the behavioural effect of adaptation, we performed a linear mixed effect (LME) model analysis, using the N200 amplitude as fixed effect and participants as random effects (PSE∼N200+(1|subjects)). The results of this analysis showed that the changes in the amplitude of the N200 could successfully predict the behavioural performance in the duration discrimination task (i.e., the PSE; R^2^ = 0.34, *β* = 2.91, t = -2.645, p < 0.001). In addition to the analysis based on the set of channels of interest chosen for the N200 component, we also applied the regression model to each of the 32 electrodes individually, in order to better localize the activity source that best predicts behavioural performance. The topography of beta values obtained by this analysis (Fig. 2C) shows that PSEs are most consistently predicted from brain activity at right occipital scalp locations (peak at channel O2; *β* = 3.5, R^2^ = 0.53, p < 0.001), contralateral to the reference stimulus position on the screen.

**FIGURE 2.**
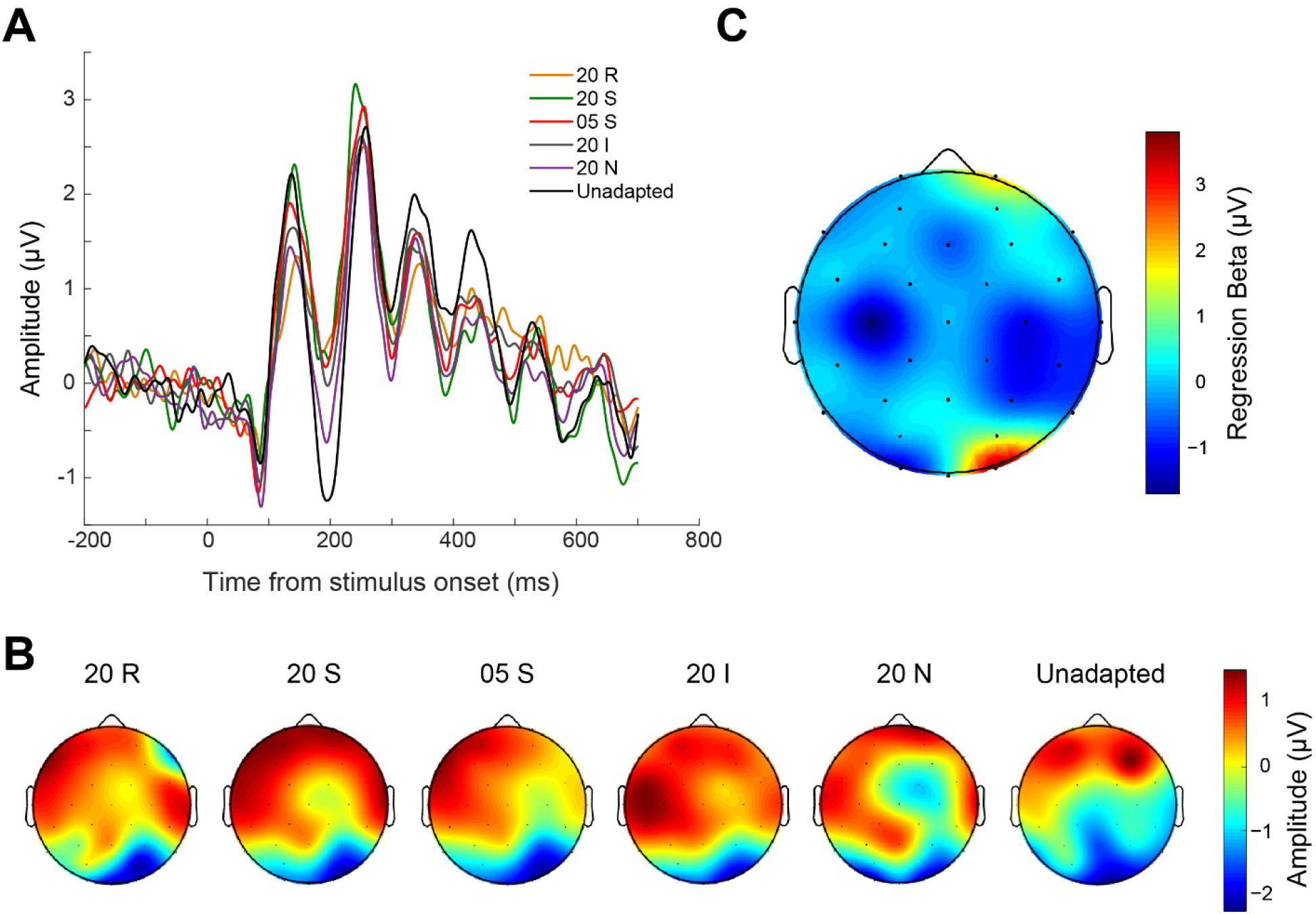
ERP results. (A) Event-related potentials (ERPs) recorded during the reference stimulus presentation (0 marks the reference onset). Each brainwave corresponds to a specific adaptation condition. (B) Scalp distribution of activity averaged across the N200 time window (140-240 ms), for each adaptation condition. (C) Topographic plot of the regression model (PSE∼N200+(1|subjects)) results (beta values).

Additionally, we also assessed whether and to what extent behavioural performance could be predicted by a late (from to 250 to 500 ms after stimulus reference onset) ERP component i.e., the contingent negative variation (CNV) component, usually observed at centro-frontal electrodes (FC1, FC2, Fz, Cz). The CNV has indeed been associated with temporal processing in previous studies (Elbert et al., 1991; Kononowicz & Penney, 2016; Li et al., 2017; Peters et al., 1977; Pfeuty et al., 2003; H. Van Rijn et al., 2011), and thus it may be involved also in this context. The CNV results are shown in Fig. S2 (see *Supplementary Online Materials*). The analysis however showed no significant difference in CNV amplitude according to the adaptation condition (one-way repeated measures ANOVA: F (4,31) = 1.09, p = 0.36), and no predictive power of CNV amplitude on the behavioural performance (LME regression, R^2^ = 0.3 average *β* = 2.36, p = 0.48).

In addition to assessing the effect of adaptation in the time domain, we also computed the impact of adaptation in the frequency domain – i.e., the effect of adaptation on oscillatory activity in the left occipito-parietal region (channels Oz, O1 and PO3) – the region where the stimuli were visually processed. Before looking at oscillatory activity during the reference stimulus presentation, we thought important in the first place to check for potential frequency entrainment during the adaptation phase and to see how this entrainment may unfold over time. Specifically, we computed changes in the power of Theta (4-7 Hz), Alpha (8-13 Hz) and Beta (13-30 Hz) frequency bands, during the adaptation phase compared to the unadapted condition. As shown in Fig. 3A, most of the frequency bands showed an increase in power compared to the unadapted state, throughout the adaptation phase, i.e., from onset to offset and extending 1 s after the adaptation phase (16 s after the adaptation onset). To better assess the changes in oscillatory activity at the offset of the adaption phase, just before the reference stimulus presentation, we took the average change in power across different frequency bands in a time window of 1 second after adaptation offset (Fig. 3B). As shown in Fig. 3B, the most prominent changes in power at the adaptation offset appeared to occur in the Beta band. A two-way repeated measure ANOVA with factors “frequency band” (i.e., Theta, Alpha, Beta) and “adaptation condition” (20N, 05S, 20I, 20S, 20R) showed a main effect of frequency band (F(2,31) = 5.2153, p = 0.0081, η^2^ = 0.0538), no main effect of adaptation condition (F(4,31) = 1.24, p = 0.23), and no interaction (F(8,31) = 0.81, p = 0.59). A series of tests collapsing together the different adaptation conditions and comparing changes in power across different frequency bands (Bonferroni-corrected α = 0.0167) showed that Beta band power changes were significantly different from both the Theta and Alpha band (t(31) = -2.6, p < 0.001, and (t(31) = -2.5, p = 0.002, respectively), while no significant difference between Theta and Alpha power changes was observed (t(31) = -0.3, p = 0.7078).

**FIGURE 3.**
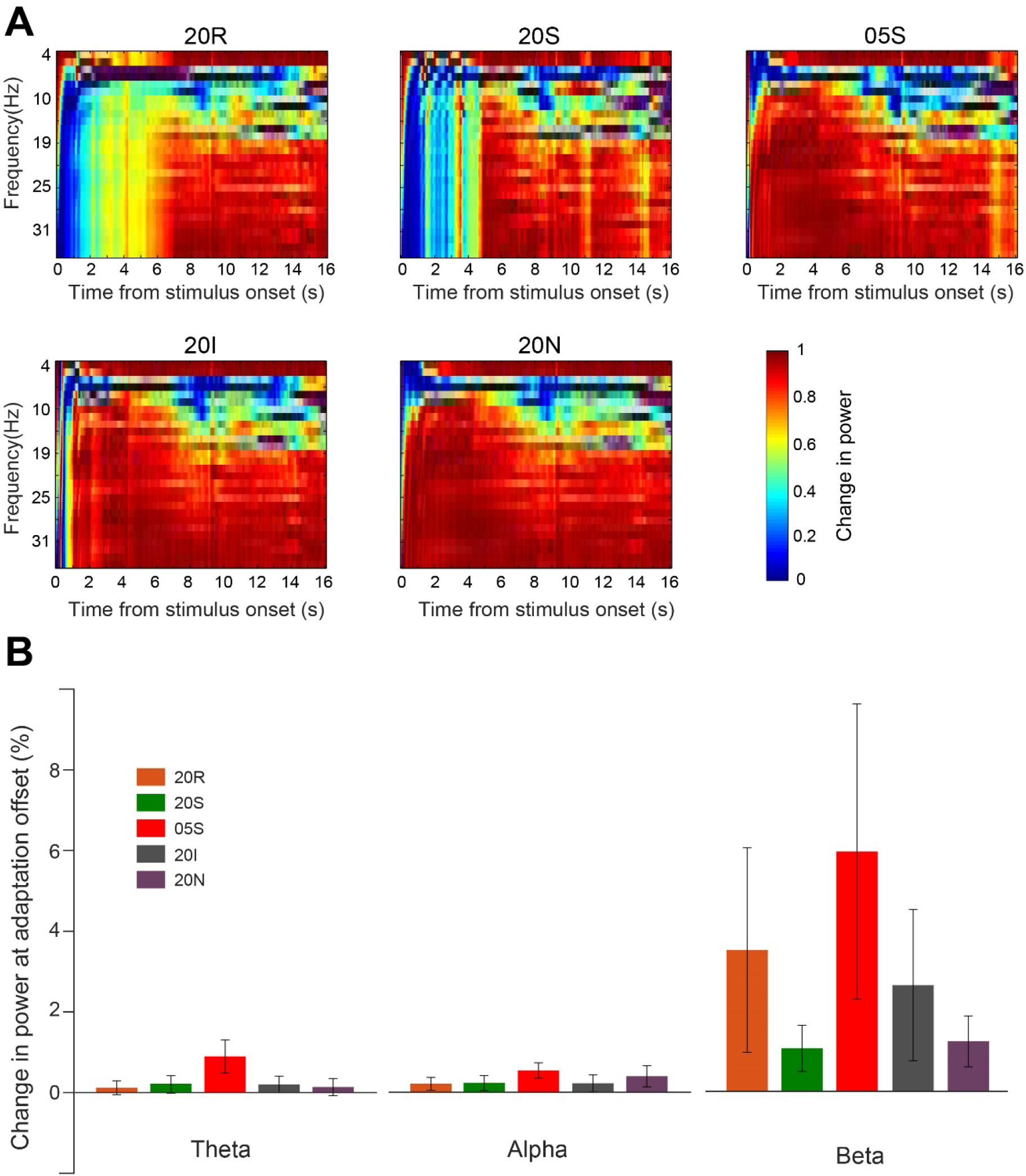
Effect of adaptation in the frequency domain. (A) Frequency changes compared to the unadapted state during the adaptation phase until its offset (adaptation phase is 15s long), across all adaptation conditions. (B) Frequency changes at the adaptation offset (1-s after adaptation offset). Effect of adaptation on oscillatory brain activity in different frequency bands (Theta = 4-7 Hz, Alpha = 8-13 Hz, Beta = 13-30 Hz) in each condition, computed at the adaptation offset. Error bars are SEM.

In order to check whether the perceptual bias caused by adaptation (Fig. 1B) was linked to the Beta band increase at the adaptation offset, we performed a LME regression analysis, using the following regression model: Adaptation effect = Beta change+adaptation condition+(1|subjects), where Beta change and adaptation condition (05S, 20S, 20I, 20N, 20R) are fixed effects and subjects represent the random effect. The results showed a good overall model fit (R^2^ = 0.35), and that the different adaptation conditions with the exception of no-topic could be predicted by changes in the Beta power (t ranging from -2.63 to -5.92, all p ≤ 0.003; no-topic condition: t = -0.88, p = 0.37). Most importantly, Beta power changes successfully predicted the perceptual bias induced by adaptation (t = 2.35, p = 0.02).

Very similar results were obtained when we looked at the adaptation effects in the 5-s top-up phase (the 5-s adaptation in each trial). As shown in Fig. 4, we observed the most robust increase in power at frequencies corresponding to the Beta band, throughout the top-up phase epoch. And similarly to the adaptation phase, also here the linear mixed-model regression analysis showed that the average change in Beta power successfully predicted the perceptual bias (*β* = 5.7, t = 1.98, p = 0.033). Also, in this model (overall model fit, R^2^ = 0.51) with the exception of no-topic adaptation (*β* = 1.78, t = -0.73, p = 0.46), Beta power changes predicted all the adaptation conditions (*β* ranging from 1,78 to 13.4, t ranging from -2.26 to -5.77, all p ≤ 0.003).

**FIGURE 4.**
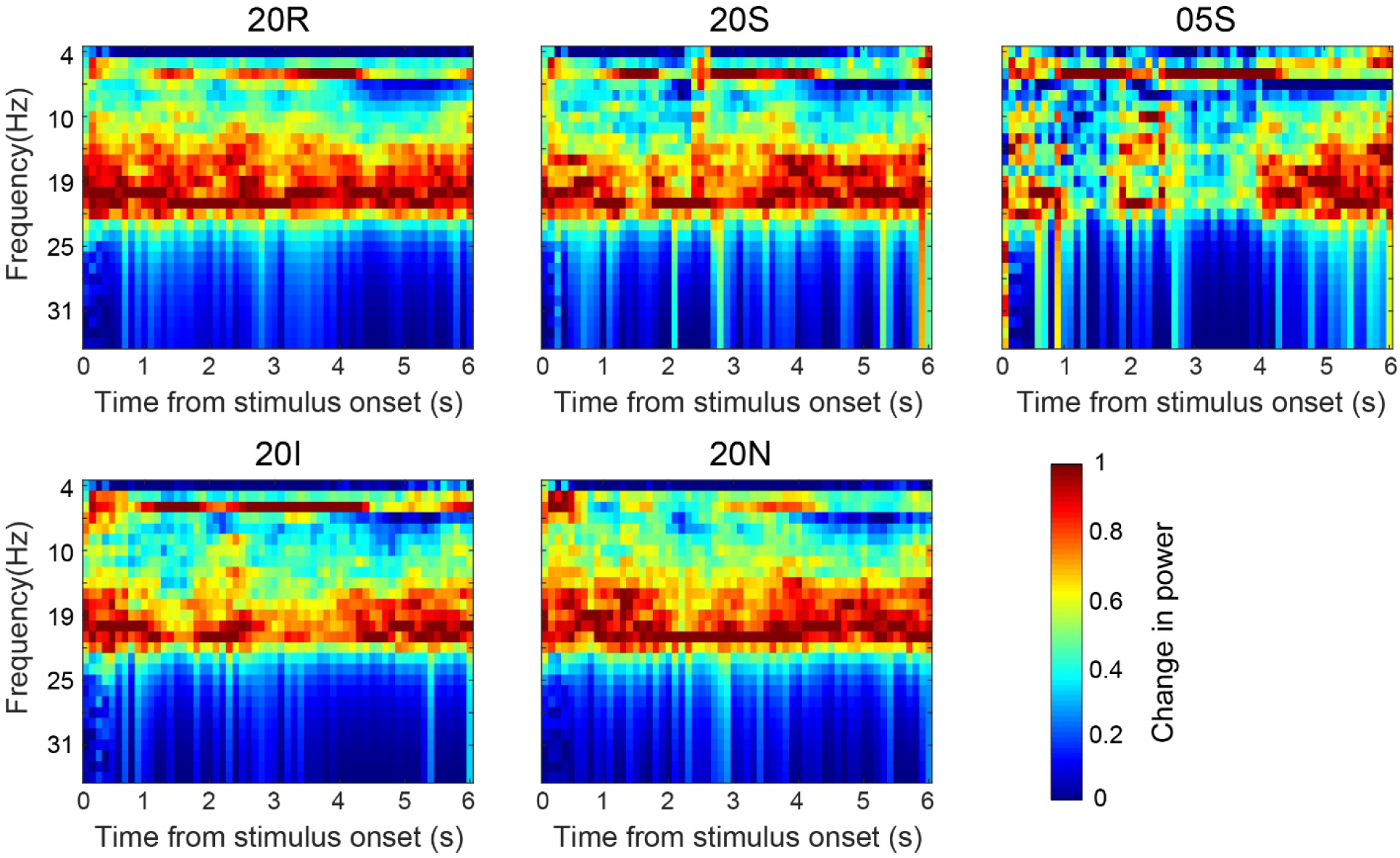
Adaptation effect in the frequency domain during the top-up phase. Changes in spectral power induced by adaptation compared to the unadapted state. The adaptation effect here was assessed during the adaptation top-up phase at the beginning of each trial across all adaptation conditions. 0 marks the top-up onset (top-up duration is 5 s)

Furthermore, since we observed a significant effect of adaptation in the Beta band both during the adaptation and the top-up phase, we also assessed whether this modulation was sustained during the subsequent presentation of the reference stimulus. As shown in Fig. 5A, the time course of spectral power changes around the presentation of the reference stimulus revealed a consistent increase in this frequency band. To further check whether the increase in Beta power changed over time and was more or less pronounced at early versus late latencies, we averaged the Beta frequency power over two distinct time windows, one spanning from 0 to 250 ms (i.e., first half of the reference stimulus duration) and the other one from 250 to 500 ms after reference onset (i.e., second half of the reference stimulus duration). As shown in Fig. 5B, we observed systematic changes in Beta power during the presentation of the first half of the reference stimulus and these effects appeared to be greatest in the adaptation conditions leading to the strongest behavioural effects (i.e., 20S and 20R). A one-way repeated measure ANOVA indeed shows a significant main effect of adaptation condition on changes in Beta power (F(4,31) = 4.3, p = 0.0236, η^2^ = 0.57). A series of one-sample t-tests (Bonferroni-corrected α = 0.01) further show that the effect was significantly greater than zero in the 20S (t(31) = 3.2, p = 0.0038) and 20R adaptation conditions (t(31) = 2.76, p = 0.0076), while no significant effect was observed in the 20I, 20N, and 05S adaptation conditions (t(31) = 1.89, p = 0.013, t(31) = 1.67, p = 0.02 and t(31) = 1.71, p=0.015 respectively). No significant difference was observed between the 20S and 20R condition (paired t-test, t(31) = 0.285, p = 0.67). In the second half of the reference presentation (250-500 ms), instead, no main effect of adaptation condition was observed (one-way repeated measure ANOVA, F (4,31) = 0.41, p = 0.8396), and changes in Beta power were not significantly greater than zero in none of the adaptation conditions (one-sample t-tests, all p > 0.05).

**FIGURE 5.**
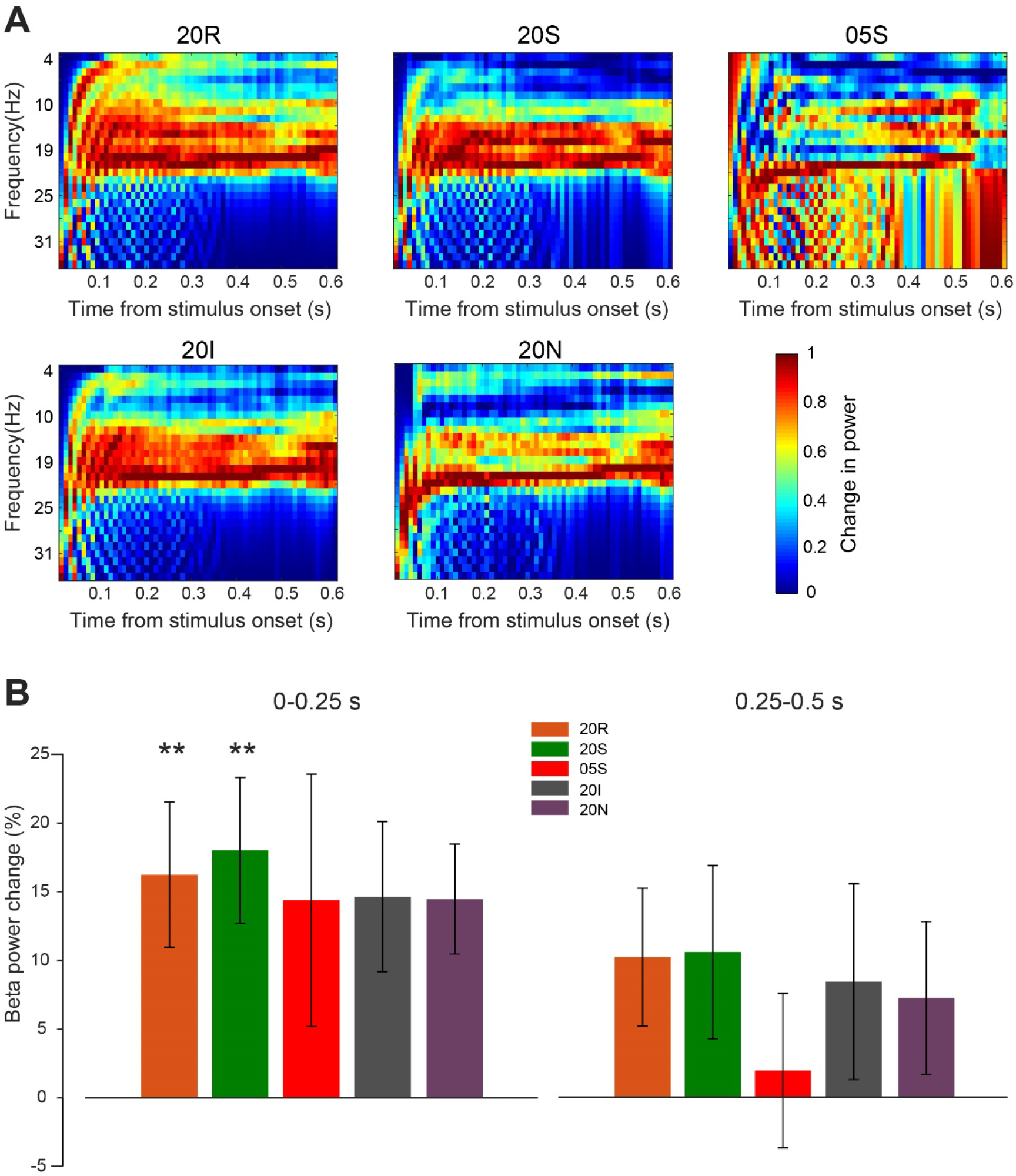
Effect of adaptation on Beta-band activity during the reference stimulus presentation. (A) Changes in spectral power in all adaptation conditions compared to the unadapted state observed during the reference stimulus presentation across multiple frequency bands. (B)Average changes in Beta power during the reference stimulus presentation in its first half (left panel: 0-0.25s after stimulus onset) and in its second half (right panel: 0.25-0.5 s). Error bars are SEM. Stars indicate the significance level of one-sample t-tests against zero ** p < 0.01.

We then assessed whether the perceptual effects induced by adaptation could be predicted by oscillatory Beta-band changes during the first half of the reference stimulus presentation (0 to 250 ms after reference onset). We performed a linear mixed effect regression analysis by fitting the following model: AE ∼ Beta(0-250ms) + adaptation condition + (1|subjects). The results (overall model fit, R^2^ = 0.37) showed that the perceptual distortions in the 20S, 20R, and 05S adaptation conditions can be successfully predicted the changes in Beta power observed in those conditions (*β* ranging from 1.46 to 12.9, t ranging from -2.00 to -5.00, all p ≤ 0.03). No significant relation between perceptual distortions and changes in Beta power was observed instead in the 20N (t = -0.42, p = 0.67) and 20I (t=-1.9 p=0.052) adaptation conditions.

Finally, Fig. 6 shows the topographic distribution of Beta power changes at the time of adaptation/top-up phase (Fig. 6A), and during the reference stimulus presentation (Fig. 6B). Looking at the scalp distribution of Beta power increase during the presentation of the adaptor stimulus, there is a clear peak at occipito-parietal scalp locations contralateral to the stimulus position (i.e., left hemisphere for the 20S, 05S, 20I, and 20N conditions, right hemisphere for the 20R condition). This suggests that the Beta power increase occurred in visual areas processing the adaptor stimulus. Interestingly though, the Beta power increase at the time of the reference presentation is instead evident, in most of the conditions (i.e., all except the 20R condition), in the opposite hemisphere compared to the effect during adaptation, at scalp locations contralateral to the reference stimulus position (see Fig. 6B). This last result suggests that the carry-over effect of adaptation is remapped to different neural populations after the gaze shift from the adaptor to the reference stimulus presentation.

**FIGURE 6.**
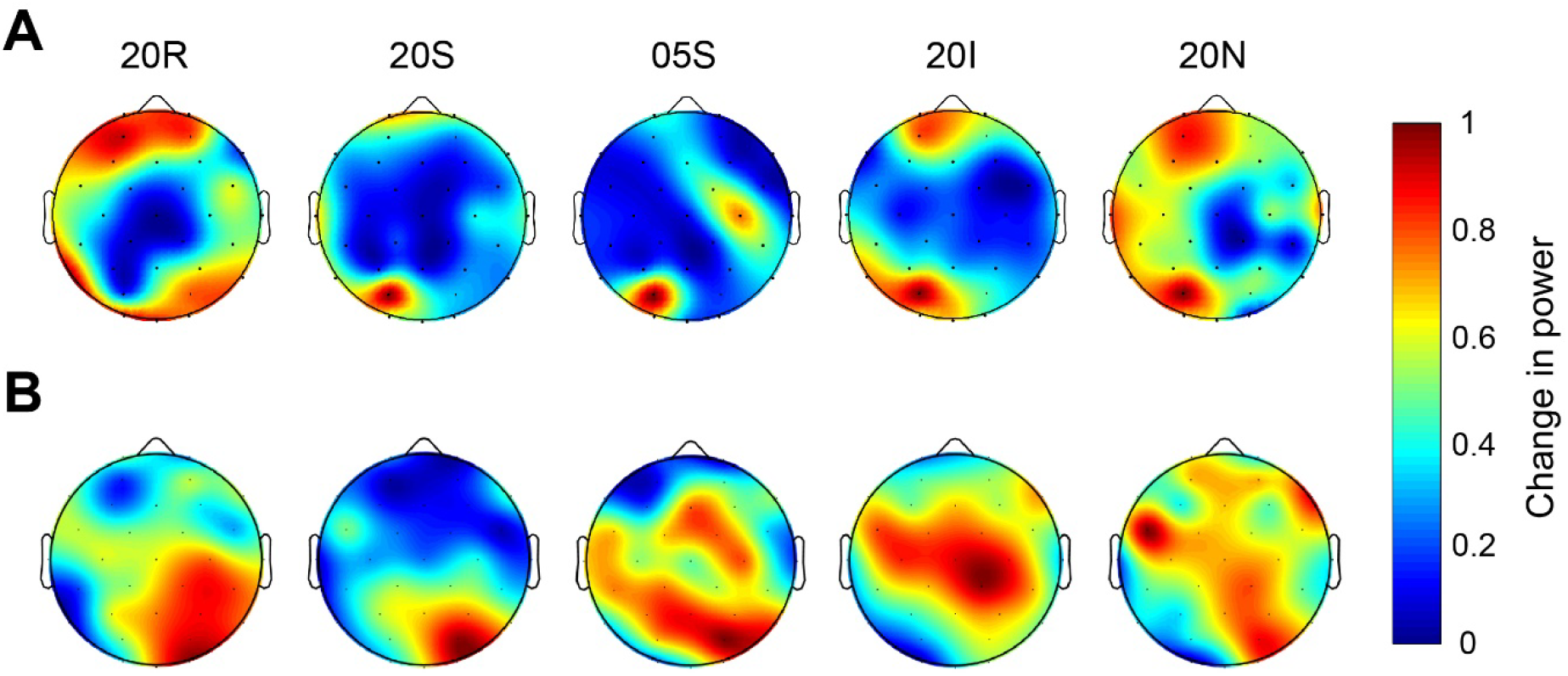
Topographic plots of Beta power changes across the scalp. **(A)** Scalp distribution of Beta power changes during the adaptation and top-up phase (collapsed together). (B) Scalp distribution of Beta power changes during the first half of the reference stimulus presentation.

Finally, we performed an exploratory decoding analysis to assess how motion adaptation affects the pattern of brain responses evoked by the reference stimulus. To control for spurious adaptation effects on brain activity not linked to the processing of the reference stimulus, in this analysis we used the no-topic adaptation condition as baseline. Thus, we basically performed a series of pairwise comparisons with the no-topic condition against all the other adaptation conditions.

Fig. 7 shows the results of the decoding analysis throughout the time course of stimulus reference processing, in terms of difference from average baseline (i.e., pre-stimulus) classification accuracy (CA). In general, the decoding performance across the different adaptation conditions showed several peaks across the reference processing time-course, suggesting that adaptation impacts multiple processing stages across the visual hierarchy. To better quantify whether and to what extent the pattern of activity evoked by the reference presented unique traces of the preceding adaptation, we binned the CA results in five different time windows spanning from 50 to 650 ms after the reference stimulus onset (i.e., 50-150 ms, 150-250 ms, 250-350 ms, 350-450 ms, and 550-650 ms). Note that due to the exploratory nature of this analysis, in the following series of t-tests p-values were corrected with a false discovery rate (FDR) procedure instead of a Bonferroni correction. In the earliest window, all adaptation conditions except one (20R; one-sample t-test, t(27) = 1.07, p = 0.29) showed a significant difference from baseline (one-sample t-tests, t(27) = 3.59, p = 0.005, *d* = 0.69, t(27) = 3.11, p = 0.008, *d* = 0.61, t(27) = 2.89, p = 0.010, *d* = 0.57, respectively for 20S, 20I, and 05S). In the second window (150-250 ms) we observed an identical pattern, with a significant decoding accuracy in the case of the 20S, 20I, and 05S adaptation condition (t(27) = 4.29, p < 0.001, *d* = 0.84, t(27) = 2.95, p = 0.013, *d* = 0.57, t(27) = 2.69, p = 0.016, *d* = 0.53, respectively; 20R: t(27) = 2.09, p = 0.046). In the intermediate window (250-350 ms), only the 20S and 20I adaptation condition showed a decoding performance significantly higher than baseline (t(27) = 3.40, p = 0.005, *d* = 0.63, t(27) = 3.34, p = 0.005, *d* = 0.64), while the 05S and 20R did not reach significance (t(27) = 2.43, p = 0.029, t(27) = 1.98, p = 0.057). At the fourth time window (350-450 ms), the difference from baseline CA appears to be generally higher compared to all other windows. All conditions in this window indeed showed a decoding performance significantly higher than baseline (20S: t(27) = 3.34, p = 0.0025, *d* = 65; 20I: t(27) = 4.22, p = 0.001, *d* = 0.80; 05S: t(27) = 3.38, p = 0.0025, *d* = 0.59; 20R: t(27) = 3.56, p = 0.025, *d* = 0.69). Finally, also at the last time window after reference offset (550-650 ms) all the condition comparisons showed a decoding performance significantly higher than baseline (20S: t(27) = 2.93, p = 0.014, *d* = 0.55; 20I: t(27) = 2.45, p = 0.021, *d* = 0.46; 05S: t(27) = 3.38, p = 0.009, *d* = 0.55; 20R: t(27) = 2.46, p = 0.021, *d* = 0.48).

**FIGURE 7.**
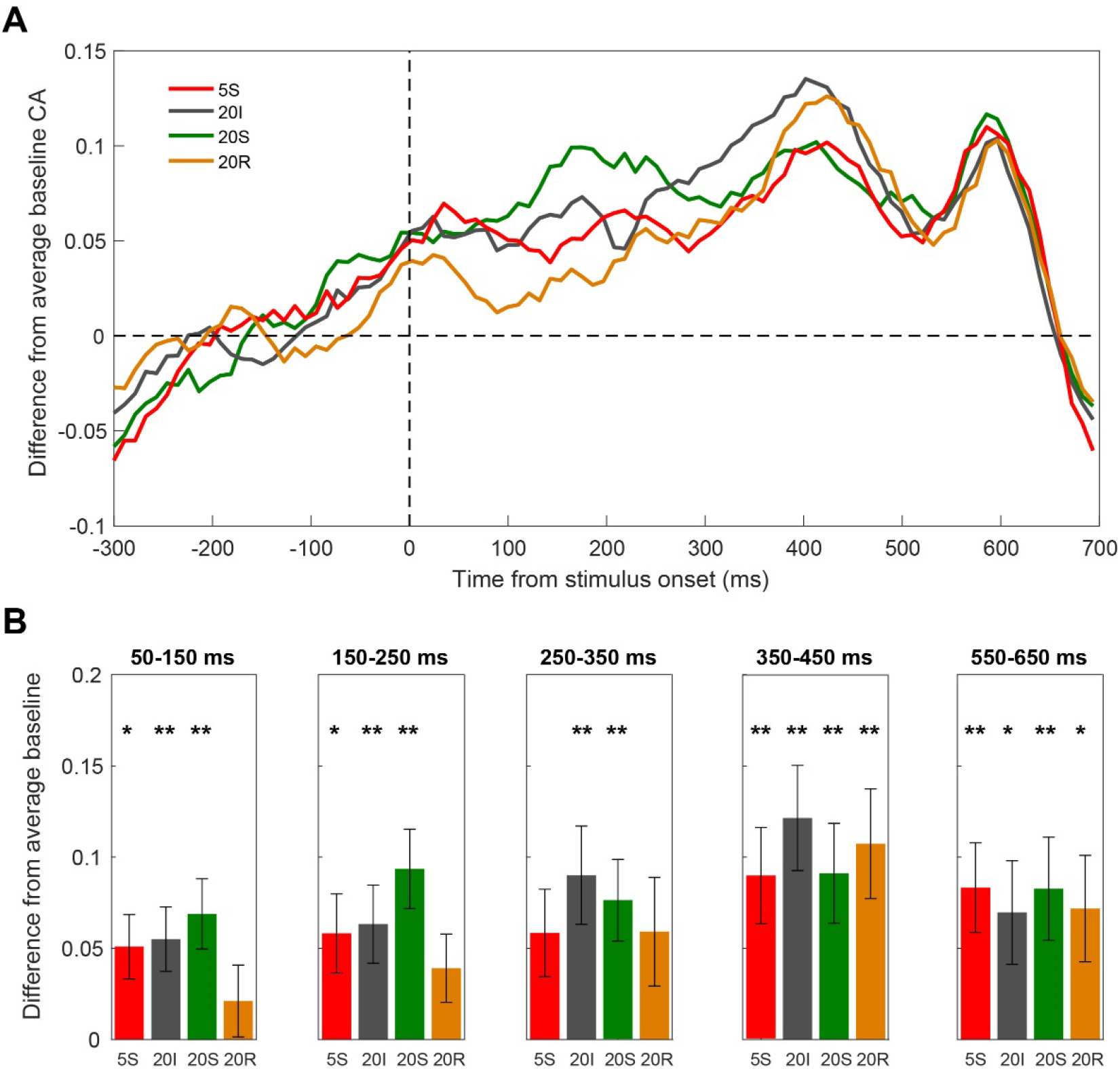
Results of the exploratory multivariate pattern analysis in the time domain. In this analysis, a pattern classifier (support vector machine) was trained on a subset of data coming from two adaptation conditions, and then tested on an independent subset of trials to predict the corresponding adaptation condition. The classification accuracy obtained in this procedure thus represents the extent to which the pattern of brain activity across multiple channels at each time point differs after different kinds of motion adaptation. This in turn highlights the different processing stages affected by adaptation, and potentially linked to perceptual time distortions. As adapting to a motion stimulus for a relatively long time is expected to leave a strong lingering trace on brain activity, we tried to minimize the possibility of spurious effects by (1) using the no-topic adaptation as a “baseline” condition (i.e., by performing pairwise comparisons of each adaptation condition against the no-topic adaptation); (2) by computing the average classification accuracy (CA) in the prestimulus interval (−300-0 ms) and subtracting it from each time point throughout the analysed epoch. This last computation was done to subtract the unspecific (i.e., not related to the reference stimulus processing) effect of different types of motion adaptation on brain activity. (A) Decoding results across the entire reference time window. Each line corresponds to the CA of each adaptation condition compared to the no-topic condition. (B) Decoding results averaged across different time windows across the post-stimulus interval. Again, the results are reported in terms of difference from average baseline CA. Asterisks refer to the significance of one-sample t-tests corrected with FDR. Error bars are SEM. * adjusted p-value < 0.025, ** adjusted p-value < 0.01

## DISCUSSION

In the present study, we investigated the neural correlates of perceptual time distortions caused by motion adaptation. Our results showed that spatiotopic and retinotopic 20 deg/s adaptation caused significant distortions in the perceived duration of the adapted reference stimulus and these distortions are coupled with changes in brain activity as measured with EEG. More specifically, we observed that modulations of the N200 amplitude time-locked to the reference stimulus presentation and recorded in occipital electrodes contralateral to it, reflect the magnitude of duration distortions measured at the behavioural level. Moreover, we also observed robust and selective increases in Beta oscillatory activity power during the adaptation/top-up phase and during the first half of the reference stimulus presentation. Crucially, we showed that such increase in Beta power predicts the perceptual duration compression effect measured behaviourally. Overall, our results thus show that duration distortions induced by motion adaptation are associated with brain changes that are (1) local – i.e., they occur in topographically-organized visual cortices processing the stimulus, and (2) they are early – i.e., as suggested by the N200 component and the Beta power modulation observed during the first half of the stimulus presentation.

Regarding the behavioural effect of adaptation, our results are in line with earlier studies showing that adaptation to a fast-moving stimulus causes a robust compression of the perceived duration of a subsequent stimulus (Ayhan et al., 2009; Bruno et al., 2010; Burr et al., 2007; Fornaciai et al., 2016; Johnston et al., 2006). In terms of the spatial reference frame of such adaptation effect, it is interesting to note that we observed an effect in both retinotopic and spatiotopic coordinates. Whether the effect of adaptation is localized in a spatial or retinal reference frame has been hotly debated in the past, due to different results supporting either interpretations (Bruno et al., 2010; Burr et al., 2011; Burr et al., 2007; Latimer et al., 2014). Such different results in turn led to interpreting the origin of adaptation as either very early (i.e., lateral geniculate nucleus or primary visual cortex) or relatively late in the visual processing stream (i.e., V5/MT). The present findings, together with other studies similarly observing adaptation in both reference frames (Fornaciai et al., 2016), suggest that adaptation may be driven by activity at multiple levels of visual processing (see Bruno & Cicchini, 2016 for a review). Our results however also show that the effect is contingent on the motion direction of adaptor and adapted stimulus, occurring only when they move in the same direction. This further suggests that a crucial contribution to the observed effect comes from motion direction-selective visual areas such as MT (Ayhan et al., 2009; Bruno et al., 2013b; Curran & Benton, 2012; Latimer et al., 2014).

Regarding the EEG results, we first show that the amplitude of the N200 ERP component is systematically modulated by adaptation at posterior scalp locations contralateral to the presented stimuli. Importantly, such modulation predicts the distortion in perceived duration observed at the behavioural level. This finding shows that the neural signature of the adaptation effect is local, and occurs in topographically organized visual areas. In this context, it is interesting to note that the N200 component has been previously linked to motion processing and adaptation (Hoffmann et al., 2001), and it has been shown to arise from the motion area V5/MT (Probst et al., 1993). The modulation at the level of the N200 component thus further supports the idea of a crucial contribution to the adaptation effect of area V5/MT.

Besides the ERP analysis, we also assessed the adaptation effect on oscillatory activity by computing changes in spectral power of oscillations in the Theta, Alpha and Beta frequency bands, compared to the non-adapted state. The results of this analysis show that the power of Beta oscillations is more substantially enhanced compared to other frequency bands, especially during the top-up phase and the first half of the reference stimulus presentation. Importantly, we show that increases in Beta power can successfully predict the perceptual bias observed at the behavioural level.

A first question thus concerns the neural mechanisms underlying such increases in Beta power and their role in temporal processing. A possibility is that an increase in power of oscillatory activity across different frequency bands might be related to the visual entrainment caused by the broadband temporal frequency of our adapting stimulus (i.e., as dot arrays have broadband spatial frequency, this results in a broad temporal frequency spectrum). However, the selective enhancement of Beta power suggests a more direct relationship between activity in this frequency band and the perceptual bias induced by adaptation. The observation that increases in Beta power can also predict perceptual distortions measured at the behavioural level further suggests an important role of Beta oscillations in temporal processing. Previous studies have indeed highlighted a role of beta-band activity in temporal encoding (Kononowicz & van Rijn, 2015; Kulashekhar et al., 2016). For instance, Kononowicz & van Rijn (2015) showed that the power of Beta activity can successfully predict the duration produced by participants in a duration reproduction task. In light of the task employed in this previous study (i.e., participants reproducing the duration by pressing a key to signal the onset and the offset of their estimated interval), changes in Beta activity were interpreted as reflecting motor inhibition processes, which in turn impact the estimated duration. Differently from this previous study, our results show that changes in Beta activity can be induced by visual adaptation, and these changes can predict the perceived duration of a visual stimulus in the absence of an explicit mapping of duration to a motor action. This reinforces the idea of an involvement of Beta activity in temporal processing independently from motor-related activity (see also Kulashekhar et al., 2016), and further suggests a role of Beta oscillations in the representation of subjective time. Differently from Kulashekhar and colleagues’ work where Beta changes were greater in fronto-central electrodes, the Beta changes observed here were mostly prominent in the occipital cortex processing the stimuli (the left hemisphere for adaptation and the right hemisphere for the reference stimulus), again suggesting a close link between local changes in brain activity and duration perception.

Besides an explicit role of Beta activity in temporal processing, an alternative idea is that Beta oscillations might reflect the maintenance of the “status quo” – i.e., the current cognitive or sensorimotor set (see Engel & Fries, 2010 for a review). Increased Beta band activity has been associated with changes in perception or with modulation of stimulus processing (Lalo et al., 2007; Okazaki et al., 2008) and this effect has been interpreted as reflecting an active maintaining of the current cognitive/sensorimotor set (Engel & Fries, 2010). In this context, the increased Beta power might reflect the change in (temporal) perceptual processing caused by adaptation, and the subsequent maintaining of this state at later latencies. In this scenario, Beta activity would thus have an indirect role in temporal processing. Although this is an interesting possibility, the finding that changes in Beta power could predict the behavioural effect (and vice versa) suggests a more direct role of oscillatory activity in carrying temporal information or modulating our subjective sense of time.

Central to our findings, is that such modulation of Beta activity occurs in a local fashion, emerging at scalp locations contralateral to the position of the stimuli. Interestingly, while the Beta power increase occurs at channels contralateral to the adaptor position, this effect shifts to the opposite hemisphere after an eye movement. Indeed, at the time of the reference presentation, the peak increase in Beta power is evident at scalp locations contralateral to the reference stimulus itself. This shift suggests that the Beta power increase is not a completely passive phenomenon, according to which we would expect an effect limited to the adapted neuronal population. But it seems instead an active process of remapping across different neuronal populations. Such a remapping may be linked to the spatial coding of the adaptation effect, that may be spatiotopic rather than retinal (e.g., see for instance Cicchini, Binda, Burr, & Morrone, 2013). However, an exception to this pattern is the retinotopic condition, where we would have expected a shift in the opposite direction (i.e., from the right to the left hemisphere) to maintain the effect in spatiotopic reference frame. The effect may thus reflect a more generalized process, occurring selectively during the adaptation phase, but then re-emerging at the time of the reference stimulus according to its position, without a stable coding in either retinotopic or spatiotopic coordinates. This may indicate a more high-level effect, possibly mediated by feedback signals from downstream areas. Assessing the specific nature of this effect however remains an open question for future investigations.

In addition to the ERP and Beta frequency results, the exploratory multivariate “decoding” analysis shows that adaptation has a strong impact on the activity evoked by the reference, affecting the pattern of brain responses throughout the entire window of reference processing. This further supports the idea that motion adaptation affects multiple stages of visual processing, and hence that the adaptation effect could be driven by activity at different stages (Bruno & Cicchini, 2016). One interesting difference in the pattern of effects of different conditions is the different time-course of the spatiotopic and retinotopic effect. Indeed, while we observed a significant effect of spatiotopic adaptation starting from the earliest time window analysed (50-150 ms), the retinotopic effect builds up more slowly and peaks at around 350-450 ms. Such a slower dynamics of the retinotopic effect suggests that it may be supported by different neural computations compared to spatiotopic adaptation. In retinotopic adaptation, part of the effect might be indirectly driven by distortions in perceived speed, resulting in a concurrent bias on perceived time (Brown, 1995; Kanai et al., 2006). In this scenario, the effect would thus arise only *after* building a representation of the stimulus speed. Although surprising, given the more high-level nature of spatiotopic processing, adaptation in a spatiotopic reference frame leads to changes in visual processing at earlier latencies. However, due to the exploratory nature of this analysis, these results have to be interpreted cautiously. Indeed, the results of the decoding procedure most likely reflect consequences of motion adaptation not uniquely related to time processing. This is also suggested by the fact that significant above-chance decoding level can be observed in most of the comparisons performed – even for conditions not resulting in a behavioural duration compression effect. Therefore, it is difficult to unambiguously relate any of the observed peaks of decoding to temporal processing.

Finally, it is interesting to note that while our data highlight the role of local and relatively low-level visual processing in subjective time distortions, it does not mean that the same pattern of effects would generalize to different experimental tasks and timing in different sensory modalities. The involvement of motion-sensitive ERPs, for instance, is most likely related to the motion stimuli used in the present study. On the other hand, in line with the idea of a widespread nature of neural timing mechanisms (Buonomano & Maass, 2009), we predict that using different stimuli would result in a different pattern of effects at the neural level, involving local and low-level brain circuits specific to the task and the sensory modality of the stimuli at hand.

To conclude, here we show that local and relatively low-level perceptual computations are critically involved in determining our subjective sense of time. Although limited to the case of motion adaptation and motion stimuli, the results support the contribution of early visual areas to the representation of time. Furthermore, they highlight the crucial role of Beta band oscillatory activity in predicting distortions of perceived time. Taken together, these results thus support the idea that time perception is deeply rooted into the sensory/perceptual processing stream, and that Beta oscillations reflect the routing of temporal information and the representation of subjective time.

## MATERIALS AND METHODS

### Subjects

The experiment was conducted on 32 healthy subjects (18 females; mean age (± SD) = 25 ± 0.37), naïve to the purpose of the experiment, none of them reporting any neurological disease. Participants gave their written informed consent before taking part in the study, and were compensated for their participation with 12 Euro/hour. The study was carried out in accordance with the Declaration of Helsinki, and was approved by the local ethics committee of the Scuola Internazionale Superiore di Studi Avanzati (SISSA). Note that the sample size of the present study was chosen to be equal to the sample included in a previous study by Kononowicz & Van Rijn (2015).

### Apparatus and stimuli

The experiment was conducted in a sound attenuated temperature-controlled room. The participants performed the experiment sitting in front of an LCD monitor (100 Hz refresh rate, resolution = 1920 x1080 pixels) where the stimuli were displayed. The stimuli used (i.e., adaptor, reference, and comparison) were arrays of black and white moving dots (number of dots = 50; 50%/50% of black and white dots) presented in a circular area with a radius of 3.25 deg, with their initial position determined randomly. Each dot had a limited lifetime, and was replaced by another, randomly positioned, dot after moving for 5 frames (50 ms). The speed of the adaptor stimuli was 20 deg/s in all of the conditions, except in the slower one where speed was 5 deg/s. The reference and comparison stimuli had a speed of 10 deg/s. The temporal frequency of all the presented stimuli was broadband around 20 Hz or 5 Hz for the adaptor, and around 10 Hz for the reference and probe stimuli, due to the broadband spectrum of spatial frequencies included in dot arrays. Motion coherence was 100% with leftward direction, except the condition where the adaptor direction was rightward. The stimuli were designed using the Psychophysics toolbox (Version 3; Kleiner, Brainard, & Pelli, 2007) in Matlab (version r2015b; The Mathworks, Inc.).

### Procedure

The experiment included 6 blocks of 70 trials, all performed within a single experimental session. In each block participants performed a duration discrimination task. With the exception of the baseline condition block, in all the rest of the blocks the duration discrimination was preceded by a perceptual adaptation phase (see below for more details). In the duration discrimination task, participants observed a reference and a comparison stimulus (always in this order), and had to report which one of the two lasted longer in time. The reference stimulus was always presented in the upper half of the screen cantered on the vertical midline, with a vertical eccentricity of 5 deg from the centre of the screen, and its duration was always 500 ms. The comparison stimulus was always presented in the lower half of the screen with the same vertical eccentricity of the reference, and its duration was pseudo randomly determined in each trial (300, 400, 500, 600 or 700 ms). To induce motion adaptation, an adaptor stimulus was presented before the reference according to different adaptation conditions based on the adaptor position, speed, and motion direction: (1) 20 deg/s motion adaptor presented in the same spatiotopic coordinates as the reference (20 deg/s spatiotopic adaptation; 20S); 20 deg/s adaptor presented in the same retinotopic coordinates of the reference (20 deg/s retinotopic adaptation; 20R); 20 deg/s spatiotopic adaptor moving in the opposite direction compared to the reference (20 deg/s inverse motion adaptation; 20I); 5 deg/s spatiotopic adaptation (5S); 20 deg/s adaptation presented in a neutral location not corresponding to either spatiotopic or retinopic coordinates of the reference (20 deg/s no-topic adaptation; 20N). Additionally, we tested a baseline condition in which no adaptor was presented. To dissociate spatiotopic and retinotopic adaptation, we asked participants to make a saccade during the interval between adaptor and reference, so that the subsequent reference position could correspond to the adaptor in terms of spatiotopic coordinates (fixed on the screen), or in terms of retinal coordinates shifted by the eye movement. The stimulation sequence in each trial was therefore as follows: the trial started with participant fixating on the first fixation point, located in the left half of the screen on the horizontal midline (horizontal eccentricity from the center of the screen = 5 deg). The adaptor was presented in one of three different positions for 15 s in the first trial of the block, and then for 5 s in the rest of the trials. After 300-450 ms from the adaptation offset, the first fixation point disappeared and a second one appeared on the right side of the screen (horizontal eccentricity from the center of the screen = 5 deg), cueing the participant to make a rightward saccade to the new fixation point (saccade amplitude = 10 deg). The reference stimulus was presented after 1 s from the onset of the second fixation point, to allow enough time to perform the saccade. Finally, the comparison stimulus was presented after 300-450 ms from the reference offset. At the end of the trial, the fixation point turned red, cueing the participant to provide a judgment, reporting which stimulus between the reference and the comparison lasted longer. Responses were provided by pressing the appropriate key (corresponding to “reference longer” or “comparison longer”) on a standard keyboard. After providing a response, the next trial started automatically after 1.3-1.45 s. The baseline condition involved a similar stimulus sequence, with the exception that no adaptor was presented before the reference. Each condition (i.e., different adaptation conditions and baseline) was performed separately in different blocks. The different blocks were separated by a break of at least 2 minutes to allow adaptation to decay. A block of trials took about 15 minutes, and the entire experimental session took about 90 minutes. See Fig. 1 for a depiction of the experimental procedure.

### EEG recording and analysis

During each condition, EEG was recorded continuously using 32 active electrodes, evenly distributed over the entire scalp (with positioning and naming conventions following a subset of the extended 10–20 system) using a BioSemi ActiveTwo system (BioSemi, Amsterdam, The Netherlands) as well as an electro-oculogram (EOG) using the set-up of Croft (Croft et al., 2005). The sampling rate was 2048 Hz. The EEG signal was re-referenced offline from the original common mode sense reference (Van Rijn, Peper, & Grimbergen, 1991) (CMS, positioned next to electrode Cz) to the average of two additional electrodes that were placed on the subject’s mastoids. The EEG signal was filtered using a 4th-order Butterworth band pass filter with range 0.5–45 Hz. Finally, the EOG signal was utilized for removing eye artefacts following the Revised Artifact-Aligned Averaging (RAAA) procedure. The EEG data corresponding to the reference stimulus was epoched from -200 to 800 ms around its onset. All epochs were baseline-corrected using the 200-ms pre-stimulus interval and were examined to remove any remaining artefact. We also extracted the EEG signal corresponding to the 15-seconds adaptation in each adaptation condition, spanning the entire adaptor presentation interval up to 1 s after its offset.

The epoched EEG signal was analysed both in the time and the frequency domain. In the time domain, we focused on EEG activity time-locked to the reference stimulus, and computed ERPs by averaging all the reference epochs corresponding to each individual baseline and adaptation condition. In order to capture visual evoked responses to the reference, we selected a set of posterior occipito-parietal channels (O1, Oz, O2, PO2, PO3), and two time windows of interest related to motion processing (N2: 140-240ms ms; (Hoffmann et al., 2001)) and to time perception (contingent negative variation, CNV: 250-500 ms; Kononowicz & Penney, 2016; Li et al., 2017; Peters et al., 1977; Van Rijn et al., 2011).

In the frequency domain, we performed a FFT analysis to compute changes in theta (4-7 Hz), alpha (8-12 Hz) and beta (15-30 Hz) frequency bands in each conditions, both considering epochs time-locked to the onset of the reference, and the initial 15-seconds period of adaptation. In the case of the reference epochs, the FFT was performed on each trial separately, and power across the different frequency bands was computed as an average of all individual trials. All the preprocessing and analytical procedures were carried out in Matlab (The Mathworks, Inc.; version r2015b).

### Behavioural data analysis

To assess the behavioural performance in the duration discrimination task, we fitted a cumulative Gaussian function to all the trials obtained in each condition (i.e., proportion “comparison stimulus longer” responses as a function of the different comparison durations), individually for each participant. We then defined the point of subjective equality (PSE), which corresponds to the perceived duration of the reference stimulus, as the median of the cumulative Gaussian function. In order to estimate the change in perceived duration induced by adaptation, we computed an adaptation effect index as the difference in PSE between each adaptation condition and the baseline unadapted condition, normalized for the baseline and turned into percentage, according to the following formula:

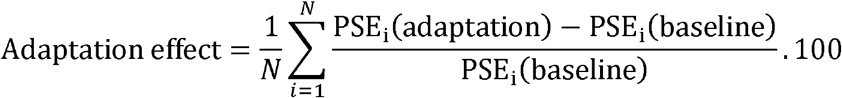

Where N is the number of subjects. The significance of the adaptation effect in different conditions was tested with a one-sample t-test against a null hypothesis of zero effect. To assess whether there was a significant difference in the effect across different conditions we performed a one-way repeated measure ANOVA with factor “adaptation condition.” Furthermore, a series of pairwise comparisons was performed, comparing each condition against each other. In all cases, the α value of the tests was corrected for multiple comparisons.

### Exploratory multivariate pattern analysis in the time domain

We also performed an exploratory multivariate “decoding” analysis (King & Dehaene, 2014) to assess how the pattern of brain activity is affected by motion adaptation, and whether and to what extent a distorted duration representation is decodable from brain signals. This technique has indeed been previously demonstrated to be sensitive to manipulations affecting a stimulus perceived magnitude (Fornaciai & Park, 2018b, 2018a, 2019). EEG data used for this analysis underwent a different preprocessing optimized to clean up the signal before the decoding procedure. First, the continuous EEG signal was epoched considering an interval from -300 to 700 ms time-locked to the onset of the reference stimulus. No baseline correction was performed in this phase of preprocessing (see below). We then used an independent component analysis (ICA) procedure to clean up the signal from eye blinks and other artefacts. After ICA, the data was band-pass filtered (0.1 to 30 Hz), and a step-like artefact rejection procedure (time window = 400 ms, step = 20 ms) was applied to further remove any residual artefact (threshold = 40 µV; average rejection rate = 2.9% ± 5.7%). Two participants were excluded following this procedure (before the analysis) due to too high rejection rates (>15% in at least one adaptation condition). Excluding such participants was necessary to allow a sufficient number of trials to perform the decoding analysis (at least 60 trials out of a total of 70 trials).

The neural decoding analysis was then performed using the Neural Decoding Toolbox (Meyers, 2013). The analysis involved a training phase where a support vector machine (SVM) classifier was trained on a subset of data obtained in specific adaptation conditions, and then in a testing phase where the classifier made predictions about which adaptation procedure was performed in the remaining subset of data. The EEG data included in the analysis was always time-locked to the adapted reference stimulus – which was identical in each condition – in order to capture the effect of adaptation on the activity evoked by the reference.

By means of this decoding procedure, we thus evaluated whether and to what extent the pattern of brain activity evoked by the reference stimulus differed as a function of the adaptation condition. More specifically, we used the no-topic condition as a “baseline,” and compared each other adaptation condition (5 Hz Spatiotopic, 20 Hz Spatiotopic, 20 Hz Retinotopic, 20 Hz Inverse motion) against it. The training and testing procedure were performed at different time windows (100 ms with a step of 15 ms) throughout the reference epoch spanning from -300 to 700 ms, with the classifier trained and tested within each individual time window. Different comparisons were tested individually, training the SVM classifier with two subsets of trials corresponding to the two conditions being compared (i.e., for example, reference activity in the no-topic vs 20 Hz Spatiotopic condition). The classifier was then tested on another subset of trials not used in the training phase (according to a leave-one-out cross validation procedure). In order to optimize the procedure, we applied a series of practices suggested by Grootswagers, Wardle, & Carlson (2017), which allow to improve the decoding procedure and improve the signal-to-noise ratio of the data included in the analysis. Namely, we first created “pseudo trials” taking the average of 10 randomly chosen trials, which provides better signal to noise ratio compared to using single trials. Second, in order to avoid overfitting, the number of EEG channels included in the analysis was limited to the five most significant ones, determined using a univariate ANOVA. Finally, this classification procedure was performed 30 times for each participant and each comparison, selecting different subsets of data for training and testing and different sets of trials to generate pseudo trials. We took the average of the 30 runs as the final estimate of the classification performance. Finally, as motion adaptation is expected to leave strong and relatively long-lasting effects on brain activity, we performed a baseline correction of classification accuracy (CA) by computing the average CA in the pre-stimulus interval and subtracting it from CA at all time points. Doing so, we thus considered the difference from average baseline CA as the net effect of adaptation on the neural activity evoked by the reference. Note that two more participants were excluded after data analysis due to anomalously high average classification accuracy across the time course of stimulus presentation (i.e., >0.85), leaving a total of 28 participants included in the reported analysis.

## Acknowledgements

This project has received funding from the European Research Council (ERC) under the European Union’s Horizon 2020 research and innovation programme -grant agreement No. 682117 BIT-ERC-2015-CoG to DB, and from the European Union’s Horizon 2020 research and innovation programme under the Marie Sklodowska-Curie grant agreement No. 838823 “NeSt” to MF.

## Author contributions

YT, BP, and DB devised the study. YT and BP collected the data. YT and MF analysed the data. YT, MF, and DB, interpreted the results and wrote the manuscript.

## REFERENCES

Ayhan, I., Bruno, A., Nishida, S., & Johnston, A. (2009). The spatial tuning of adaptation-based time compression. Journal of Vision, 9(11), 1–12. https://doi.org/10.1167/9.11.2

Brown, S. W. (1995). Time, change, and motion: The effects of stimulus movement on temporal perception. Perception & Psychophysics, 57, 105–116.

Bruno, A., Ayhan, I., & Johnston, A. (2010). Retinotopic adaptation-based visual duration compression. Journal of Vision, 10(10), 30.

Bruno, A., & Cicchini, G. M. (2016). Multiple channels of visual time perception. Current Opinion in Behavioral Sciences, 8, 131–139.

Bruno, A., Ng, E., & Johnston, A. (2013). Motion-direction specificity for adaptation-induced duration compression depends on temporal frequency. Journal of Vision, 13(12), 19. https://doi.org/10.1167/13.12.19

Buonomano, D. V, & Maass, W. (2009). State-dependent computations: spatiotemporal processing in cortical networks. Nature Reviews Neuroscience, 10(2), 113–125.

Burr, D. C., Cicchini, G. M., Arrighi, R., & Morrone, M. C. (2011). Spatiotopic selectivity of adaptation-based compression of event duration. Journal of Vision, 11(2), 21; author reply 21a. https://doi.org/10.1167/11.2.21

Burr, D., Tozzi, A., & Morrone, M. C. (2007). Neural mechanisms for timing visual events are spatially selective in real-world coordinates. Nature Neuroscience, 10(4), 423.

Cicchini, G. M., Binda, P., Burr, D. C., & Morrone, M. C. (2013). Transient spatiotopic integration across saccadic eye movements mediates visual stability. Journal of Neurophysiology, 109(4), 1117–1125.

Croft, R. J., Chandler, J. S., Barry, R. J., Cooper, N. R., & Clarke, A. R. (2005). EOG correction: a comparison of four methods. Psychophysiology, 42(1), 16–24.

Curran, W., & Benton, C. P. (2012). The many directions of time. Cognition, 122(2), 252–257.

Elbert, T., Ulrich, R., Rockstroh, B., & Lutzenberger, W. (1991). The processing of temporal intervals reflected by CNV□like brain potentials. Psychophysiology, 28(6), 648–655.

Engel, A. K., & Fries, P. (2010). Beta-band oscillations-signalling the status quo? Current Opinion in Neurobiology, 20(2), 156–165. https://doi.org/10.1016/j.conb.2010.02.015

Fornaciai, M., & Park, J. (2018a). Early numerosity encoding in visual cortex is not sufficient for the representation of numerical magnitude. Journal of Cognitive Neuroscience, 30(12). https://doi.org/10.1162/jocn_a_01320

Fornaciai, M., & Park, J. (2018b). Attractive Serial Dependence in the Absence of an Explicit Task. Psychological Science, 29(3), 437–446. https://doi.org/10.1177/0956797617737385

Fornaciai, M., & Park, J. (2019). Neural Dynamics of Serial Dependence in Numerosity Perception. Journal of Cognitive Neuroscience, 32(1), 141–154. https://doi.org/10.1162/jocn_a_01474

Fornaciai, Michele, Arrighi, R., & Burr, D. C. (2016). Adaptation-Induced Compression of Event Time Occurs only for Translational Motion. Scientific Reports, 6(August 2015), 1–13. https://doi.org/10.1038/srep23341

Fornaciai, Michele, Markouli, E., & Di Luca, M. (2018). Modality-specific temporal constraints for state-dependent interval timing. Scientific Reports, 8(1), 1–10.

Grootswagers, T., Wardle, S. G., & Carlson, T. A. (2017). Decoding dynamic brain patterns from evoked responses: A tutorial on multivariate pattern analysis applied to time series neuroimaging data. Journal of Cognitive Neuroscience, 29(4), 677–697.

Hoffmann, M. B., Unsöld, A. S., & Bach, M. (2001). Directional tuning of human motion adaptation as reflected by the motion VEP. Vision Research, 41(17), 2187–2194.

Javadi, A. H., & Aichelburg, C. (2012). When time and numerosity interfere: the longer the more, and the more the longer. PloS One, 7(7), e41496.

Johnston, A., Arnold, D. H., & Nishida, S. (2006). Spatially localized distortions of event time. Current Biology, 16(5), 472–479.

Kanai, R., Paffen, C. L. E., Hogendoorn, H., & Verstraten, F. A. J. (2006). Time dilation in dynamic visual display. Journal of Vision, 6(12), 8.

Karmarkar, U. R., & Buonomano, D. V. (2007). Timing in the absence of clocks: encoding time in neural network states. Neuron, 53(3), 427–438.

King, J.-R., & Dehaene, S. (2014). Characterizing the dynamics of mental representations: the temporal generalization method. Trends in Cognitive Sciences, 18(4), 203–210.

Kleiner, M., Brainard, D., & Pelli, D. (2007). What’s new in Psychtoolbox-3?

Kononowicz, T. W., & Penney, T. B. (2016). The contingent negative variation (CNV): Timing isn’t everything. Current Opinion in Behavioral Sciences, 8, 231–237.

Kononowicz, T. W., & van Rijn, H. (2015). Single trial beta oscillations index time estimation. Neuropsychologia, 75, 381–389.

Kristjánsson, Á., Vuilleumier, P., Schwartz, S., Macaluso, E., & Driver, J. (2007). Neural Basis for Priming of Pop-Out during Visual Search Revealed with fMRI. Cerebral Cortex, 17(7), 1612–1624. https://doi.org/10.1093/cercor/bhl072

Kulashekhar, S., Pekkola, J., Palva, J. M., & Palva, S. (2016). The role of cortical beta oscillations in time estimation. Human Brain Mapping, 37(9), 3262–3281.

Lalo, E., Gilbertson, T., Doyle, L., Di Lazzaro, V., Cioni, B., & Brown, P. (2007). Phasic increases in cortical beta activity are associated with alterations in sensory processing in the human. Experimental Brain Research, 177(1), 137–145.

Latimer, K., Curran, W., & Benton, C. P. (2014). Direction-contingent duration compression is primarily retinotopic. Vision Research, 105, 47–52.

Li, B., Chen, Y., Xiao, L., Liu, P., & Huang, X. (2017). Duration adaptation modulates EEG correlates of subsequent temporal encoding. NeuroImage, 147(2), 143–151. https://doi.org/10.1016/j.neuroimage.2016.12.015

Meck, W. H., & Benson, A. M. (2002). Dissecting the brain’s internal clock: how frontal–striatal circuitry keeps time and shifts attention. Brain and Cognition, 48(1), 195–211.

Meyers, E. (2013). The neural decoding toolbox. Frontiers in Neuroinformatics, 7, 8.

Okazaki, M., Kaneko, Y., Yumoto, M., & Arima, K. (2008). Perceptual change in response to a bistable picture increases neuromagnetic beta-band activities. Neuroscience Research, 61(3), 319–328.

Peters, J. F., Billinger, T. W., & Knott, J. R. (1977). Event related potentials of brain (CNV and P300) in a paired associate learning paradigm. Psychophysiology, 14(6), 579–585.

Pfeuty, M., Ragot, R., & Pouthas, V. (2003). Processes involved in tempo perception: a CNV analysis. Psychophysiology, 40(1), 69–76.

Probst, T., Plendl, H., Paulus, W., Wist, E. R., & Scherg, M. (1993). Identification of the visual motion area (area V5) in the human brain by dipole source analysis. Experimental Brain Research, 93(2), 345–351.

Spencer, R. M. C., Karmarkar, U., & Ivry, R. B. (2009). Evaluating dedicated and intrinsic models of temporal encoding by varying context. Philosophical Transactions of the Royal Society B: Biological Sciences, 364(1525), 1853–1863.

Togoli, I., Fornaciai, M., & Bueti, D. (2020). The functional properties of human magnitude integration. PsyArXiv. https://doi.org/10.31234/osf.io/g7w3q

Treisman, M., Faulkner, A., Naish, P. L. N., & Brogan, D. (1990). The internal clock: Evidence for a temporal oscillator underlying time perception with some estimates of its characteristic frequency. Perception, 19(6), 705–742.

Van Rijn, A. C. M., Peper, A., & Grimbergen, C. A. (1991). High-quality recording of bioelectric events. Medical and Biological Engineering and Computing, 29(4), 433–440.

Van Rijn, H., Kononowicz, T. W., Meck, W. H., Ng, K. K., & Penney, T. B. (2011). Contingent negative variation and its relation to time estimation: a theoretical evaluation. Frontiers in Integrative Neuroscience, 5, 91.

Wiener, M., Matell, M. S., & Coslett, H. B. (2011). Multiple Mechanisms for Temporal Processing. Frontiers in Integrative Neuroscience, 5. https://doi.org/10.3389/fnint.2011.00031

Xuan, B., Zhang, D., He, S., & Chen, X. (2007). Larger stimuli are judged to last longer. Journal of Vision, 7(10), 2.

