## Supplementary materials for "Subjective time is predicted by local and early visual processing"

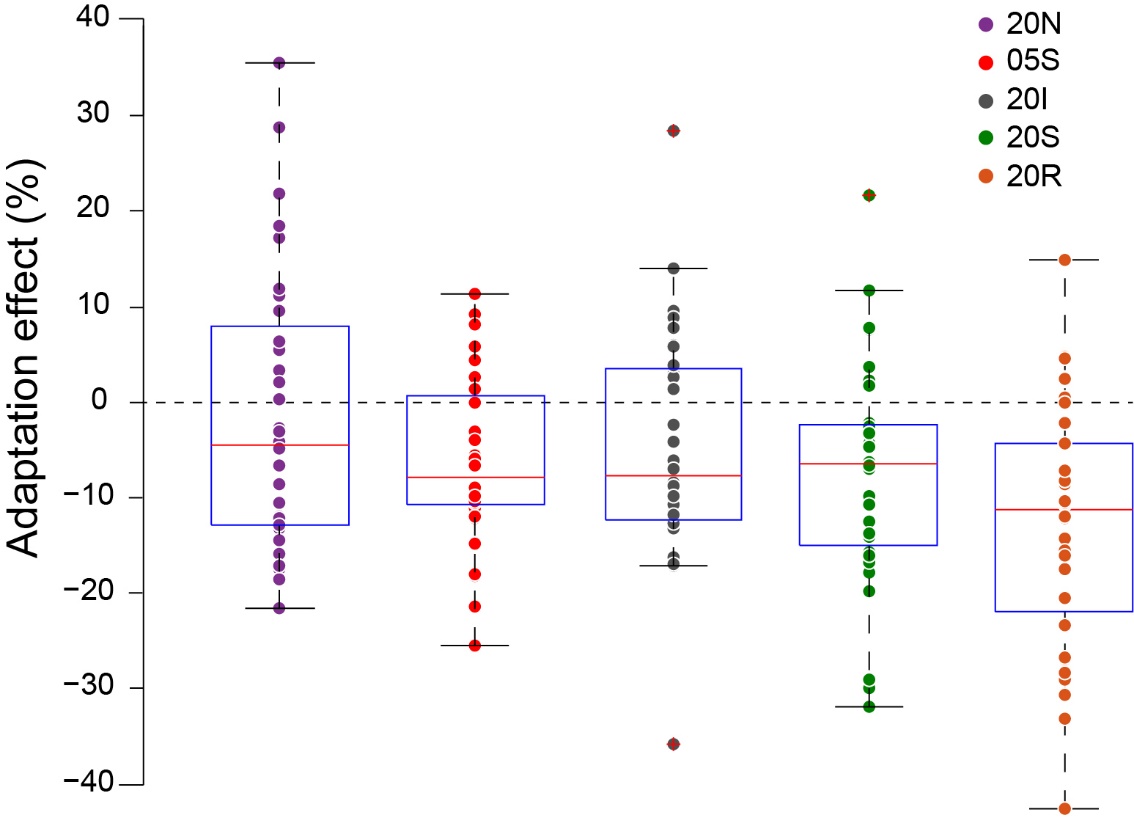

**Figure S1. Behavioural effect of adaptation**. The figure shows the overall group statistics as Box plots, while the individual effects are shown by the individual data points superimposed to the plots. The figure legend indicates the different conditions tested: 20N: 20 deg/s no-topic adaptation; 05S: 5 deg/s spatiotopic adaptation; 20I: 20 deg/s spatiotopic inverse motion adaptation; 20S: 20 deg/s spatiotopic adaptation; 20 deg/s retinotopic adaptation. Negative adaptation effect indexes indicate a reduction in perceived duration after adaptation.

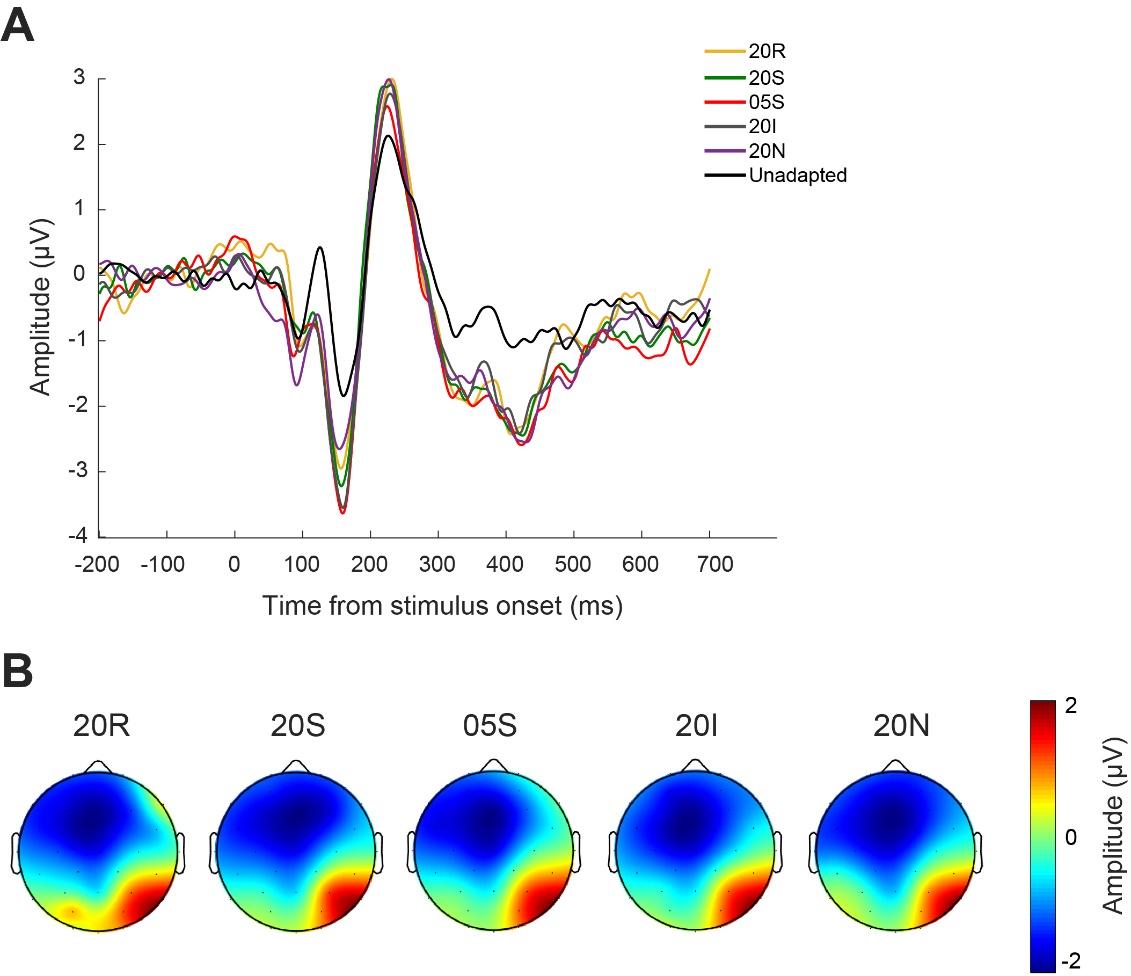

**Figure S2. CNV component**. (A) Event-related potentials time-locked to the reference stimulus onset, corresponding to the different adaptation conditions, averaged across a set of fronto-central channels where the CNV is expected to peak (FC1, FC2, Fz, Cz). (B) Scalp distribution of activity averaged across a time window corresponding to the typical timing of the CNV component (250-500 ms after stimulus onset).
